# Running System of Flipped Teaching Based on Video Conference

**DOI:** 10.1101/2022.10.10.511270

**Authors:** Xiao-Yu Zhang

## Abstract

**Objective:** I aimed to provide a basis for the standardized operation of Flipped Teaching based on Video Conference for reference by other institutions or organizations.

**Methods:** The teaching and research materials of Flipped Teaching with Video Conference as carrier carried out in April and June 2022 were collected. Based on the “Structure-Process-Outcome” theory, the induction method and SWOT analysis method were used for systematic analysis.

**Results:** A total of 43 residents participated in the teaching project, and 40 of them passed the examination, accounting for 93.0%. And the teaching project had been analyzed and reported in 3 literature. For the successful operation of this project, the following five dimensions were summarized. First, the preparatory dimension of teaching: clear teaching form, objectives, content, reference materials and assignments; Second, the carrier of teaching implementation: the selection, function and debugging of teaching equipment; Third, the teaching management dimension: the management mode, management team, lecturers team, trainees management and emergency response; Fourth, the teaching evaluation dimension: trainees’ systematic evaluation of the teaching; Fifth, the teaching assessment dimension: attendance rate, assignments completion rate and exam pass rate.

**Conclusion:** The core of the running system is to carry out Flipped Teaching with Video Conference as carrier to realize the standardized training for residents, and ensure the smooth progress and the quality of teaching through teaching management, evaluation and assessment.

## Introduction

Due to the COVID-19 pandemic, Shanghai had implemented regional control measures since March 28, 2022[1], and many internal medicine residents were unable to conduct infectious diseases training across the hospitals in April. Therefore, I explored a novel teaching model, and I tried Flipped Teaching based on Video Conference for those internal medicine residents in the designated hospital for infectious diseases training from April 1 to April 4.[3] The consequence was that the teaching model was generally effective in the infectious diseases training, so I continued this training model throughout the whole April for those residents. For further verifying the training effect and repeatability of this teaching model, I carried out this teaching model for another group of residents for infectious diseases training in June, and the study once more illustrated that this teaching model was generally effective for residents training, with good feedback and strong feasibility.[4]

In order to fully evaluate the effectiveness of the teaching model and the requirements of the resident process assessment, I conducted an online infectious disease examination for the above residents on July 11, and 93.0% passed the examination.[5] This once again confirmed effectiveness and the applicable value of this teaching model.

In view of the trial implementation in April and the verification of this teaching mode in June, and the examination results of the above residents on July 11, this teaching mode had obvious applicable value and prospects.[3-5] So, I summarized teaching and research materials of Flipped Teaching based on Video Conference both April and June to create a running system that could be referenced by other organizations or institutions.

## Objects and Methods

### Subjects

The subjects of this study were the teaching data, examination data and research literature of Shanghai residents who participated in Flipped Teaching with Video Conference as carrier in April and June 2022. And this study was conducted in accordance with the Declaration of Helsinki.

### Definition of Flipped Teaching Based on Video Conference

This teaching mode first needed to establish management and lecturers team, and selected the carrier of Video Conference. According to the requirements of “Content and Standard of Standardized Training for residents”, lecturers chose the training topics and contents, and determined the teaching time within the training cycle. After the training content and teaching time were settled, the teaching plan and implementation process for the period were formulated. Residents conducted clinical training according to the latest clinical guidelines and expert consensus, and lecturers developed PPT and clinical teaching. The whole teaching activities were carried out online by Video Conference, involving teaching organization, implementation, discussion after teaching, assignments, evaluation, assessment, examination and management.[3-5]

### Research Method

Research Theory: This research was based on the “Structure-Process-Outcome” theory.[2] The running system came from the analysis and summary of the teaching structure, process and outcome of Flipped Teaching based on Video Conference.

Induction analysis: The items in the structure design, implementation process and assessment results of the teaching were categorized and summarized.

SWOT analysis: SWOT analysis was carried out on the classified summary items of the teaching, and the strengths, weaknesses, opportunities and threats of each item were identified, subsequently conclusions were drawn.

## Results

### Basic Information of Residents and Literature

There were 19 residents in April and 24 residents in June joining the Flipped Teaching based on Video Conference for infectious disease training in the study, all from tertiary hospitals. Among them, the average age was 29.0 years old, males accounted for 34.9 %, 37.2% had a bachelor’s degree, 18.6% had a master’s degree, and 44.2% had a doctoral degree, residents in the first year of standardized training accounted for 4.7%, residents in the second year accounted for 67.4 %, residents in the third year accounted for 27.9 %, and 93.0% had passed the examination.[5] The detailed information of residents is shown in **Table 1**.

**Table 1.** Baseline of Residents Participating in Flipped Teaching

| Indicators | April(n=19) | June(n=24) |
| --- | --- | --- |
| Sexuality [male,n,(%)] | 9(47.4) | 6(25.0) |
| Age [years,median,(interquartile range)] | 30.0(27.0-32.0) | 28.5(25.3-30.8) |
| Education background |  |  |
| Bachelor's degree [n,(%)] | 5(26.3) | 11(45.8) |
| Master's degree [n,(%)] | 4(21.1) | 4(16.7) |
| Doctoral degree [n,(%)] | 10(52.6) | 9(37.5) |
| Training phase |  |  |
| First year of training [n,(%)] | 2(10.5) | 0(0) |
| Second year of training [n,(%)] | 12(63.2) | 17(70.8) |
| Third year of training [n,(%)] | 5(26.3) | 7(29.2) |
| From tertiary hospitals [n,(%)] | 19(100) | 24(100) |
| Examination score [years,median,(interquartile range)] | 72.0(63.0-78.0) | 71.0(64.3-80.0) |
| Number of examination score $\geq 60$ / Number of passing examination [n,(%)] | 18(94.7) | 22(91.7) |

There were three research reports on Flipped Teaching with Video Conference as carrier in 2022. The detailed information of literature is shown in **Table 2**.

**Table 2.** Related Literature of Flipped Teaching Based on Video Conference

| Date | Title |
| --- | --- |
| 2022/8/13 | Correlation Analysis of Demographic Characteristics of Internal Medicine Residents on the Training Effect of Flipped Teaching Based on Video Conference[5] |
| 2022/7/20 | Promotion and Validation of Flipped Teaching Based on Video Conference in Standardized Training for Internal Medicine Residents[4] |
| 2022/4/17 | Application and Evaluation of Flipped Teaching Based on Video Conference in Standardized Training for Internal Medicine Residents[3] |

### Running System

The running system included five parts: the preparatory dimension of teaching, carrier of teaching implementation, management dimension, evaluation dimension and assessment dimension.[3-6]

#### (1) Preliminary preparation for teaching

The teaching form and objectives needed to be defined, according to the requirements of the syllabus to determine the teaching content and reference materials. Next a training plan should be formulated, involving teaching organization, training, after-class discussion, quality control and teaching management. It was necessary to make requirements for teaching PPT, assignments, attendance, evaluation and assessment.

#### (2) Teaching implementation carrier

It was necessary to select, function and debug teaching equipment. The equipment had audio, video and online communication functions. The exam equipment should have the function of online examination, and at least one other equipment should be enabled for the invigilation of the online exam.

#### (3) Teaching management

The management mode adopted vertical management mode to ensure the smooth development of management and teaching; The management team was involved in the whole process of teaching management, developing and implementing teaching plans, as well as online management and supervision. The lecturers team should prepare the teaching content, perform the teaching work, and answer trainees’ questions as well. All residents needed to participate in the learning, complete the teaching contents, and ask questions about the teaching contents.

#### (4) Teaching evaluation

Six teaching plan indicators: teaching on the planned time, teaching on the planned content, making PPT fully, providing references, unifying the teaching content and training program, and participating in after-class discussion. Nine teaching quality indicators: rigorous teaching attitude, punctual class, detailed and accurate teaching content, reasonable structure and clear process, highlighting teaching key points, clear teaching difficulties, accurate and refined language, combining theory with clinical practice, improving ability to analyze and deal with the disease.[3,4]

#### (5) Teaching assessment

The teaching assessment included attendance rate, teaching discussion, assignments completion rate and examination pass rate. Teaching video was used to dynamically monitor the attendance of residents in teaching activities. All residents would discuss the teaching content, and the lectures should give answers to all the questions raised and set questions for trainees to practice. Finally, all residents were tested online with standardized questions to assess teaching quality and effect.

## Discussion

This study was a retrospective study. It was based on the theory of “Structure-Process-Outcome”, the teaching materials and research both April and June were summarized and classified, SWOT analysis method was used to further analyze, and the running system of Flipped Teaching based on Video Conference was established for other organizations or institutions to apply in the field of the standardized training of residents. To this end, I carried on this study, and following four aspects of analysis and elaboration were shown for discussion to this research.

### Advantages of this Running System

This running system was not limited by space and rarely constrained by time. It gave full play to the initiative of residents, and Flipped Teaching with Video Conference as carrier could be used as a supplementary form of standardized training for residents in special periods.[3-4]

### Deficiencies of this Running System

This running system had been applied and verified in the training of infectious diseases for internal medicine residents, and had not been verified in other fields. The running system had achieved favorable results in a single application of about 20 residents each teaching, but the effect was uncertain in terms of fewer or more people.

### Key Points of this Running System

The running system was managed by vertical management mode[6], and the responsibilities and tasks of managers, lecturers and trainees were clarified to ensure the smooth progress of teaching. Teaching evaluation and assessment were needed to ensure the quality of teaching. Focus on: the knowledge requirements of the organizers and the whole process of teaching intervention, support from superior authorities and joint training hospitals, the selection of reference materials, asking questions by all trainees and giving answers by the lecturers in the class, the lecturers setting questions according to the teaching contents and the trainees completing assignments after class, the selection and testing of teaching software, standardized question bank with the examination, perfect examination equipment, invigilation equipment and team.

### Running System Needed to be Further Optimized

This running system needs to be further studied in other professional fields of the Standardized Training of Residents. At the same time, more experience is needed in clinical operation.

## Conclusions

The core of this running system is to carry out Flipped Teaching with Video Conference as carrier to achieve standardized and homogeneous training objectives for residents, and to ensure the smooth progress and the quality of teaching through teaching management, evaluation and assessment.

## Ethical Approval and Consent to Participate

Informed consents of participants in the standardized training for those residents were obtained for the training and the study. The study received Institutional Review Board (IRB) approval by the Shanghai Public Health Clinical Center Ethics Committee. The IRB number was No. 2021-S026-01.

## Acknowledgments

This study was supported by the internal medicine residents who had participated in the standardized training for residents in Shanghai. Thanks to the teaching administration departments and the resident standardized training bases of the united training hospital for their supports.

## Authors’ Contributions

Xiao-Yu Zhang made conception, design, acquisition of data, analysis and interpretation of data, drafted and revised the manuscript, and agreed to be accountable for all aspects of the work.

## Consent for publication

The author has read and agreed to the published version of the manuscript.

## Literature share statement

The study included in the manuscript submitted to the journal is transparent. Consent from the corresponding author is required for any institution or individual to reprint this document.

